# Optimizing Disease Surveillance with Blockchain

**DOI:** 10.1101/278473

**Authors:** Flávio Codeço Coelho

## Abstract

Disease surveillance, especially for infectious diseases, is a complex and inefficient process. Here we propose an optimized, blockchain-based monitoring and reporting process which can achieve all the desired features of an ideal surveillance system while maintaining costs down and being transparent and robust. We describe the technical specifications of such a solution and discuss possibilities for its implementation. Finally, the impact of the adoption of distributed ledger technology for disease surveillance is discussed.

## Introduction

Technologies commonly known as blockchains are implementations of distributed ledgers for the safekeeping of information, in a manner which does away with the need for centralized management, and as a consequence, are censorship resistant. These Technologies are finding a growing number of applications, since the first widespread application of a distributed ledger, Bitcoin, was proposed in 2008(Nakamoto, 2008).

Blockchain technology has started to attract the attention in health management circles. An obvious application is the management of electronic health records (Ekblaw et al., 2016). Peterson et al. (Peterson et al., 2016) Argue that blockchains can solve all healthcare data exchange problems.

Disease surveillance, especially for infectious diseases, is a complex and inefficient process. Surveillance systems are typically composed of multiple stages, each focuses on a different level of Information management and analysis(Paquet et al., 2006). In this paper we will focus only on the problems of case monitoring, which is the first stage in any surveillance system.

Disease monitoring systems must aggregate data coming from a large network of agents which must report disease cases according to a pre-established protocol. The data received must be validated and then made available to health officials to help manage their response to public health demands. Due to the sensitivity of the information, These large databases are subject to central control which is usually where all chronic problems of disease surveillance systems come from. Among the main problems brought about by central control we can list: a single point of failure; delays in the aggregation of information; lack of transparency; lack of responsiveness to failures; to cite but a few.

Instead of going through the long list of shortcomings of current surveillance information systems, we can instead try to elicit the desiderata(CDC, 1988) for an ideal surveillance system. As we are focusing only on the case reporting stage, the key fetures we envision are the following.

**Provenance:** origin of a case report must always be known. This means effective geographical location but also the ID of the health establishment and/or professional collecting the data.
**Timing:** The date and time of reporting must be accurately known. This timestamp is not the same as the date of the onset of symptoms, but knowing it allows for the assessment of the readiness of of health services.
**Uniqueness:** Each case report must be uniquely identifiable, to allow for effective data management, avoiding duplicates of conflicting co-existing versions of the same case report.
**Selective privacy:** Some informations on the case report must remain private, since they consist of PII^1^. but other fields can be publicized. Privacy issues should not stand in the way of accessibility of the public information.
**Queryability:** Data must be queryable in an expressive way, i. e., allowing for filtering, grouping, ordering, etc.
**Coverage:** The network of health professionals reporting on cases, must maximize capillarity and be cheaply expandable.
**Incentives:** Health professionals must be incentivized to report promptly and accurately.
**Consensus:** Data must be validated so there is no disagreement about a report. The validation mechanism must also guarantee the veracity of the data as much as possible.
**Versioning:** Case reports are usually revised. There must be an easy way to update reports without losing original data.
**Durability:** The possibilities of data loss must be minimal. Long term retention of data must be guaranteed.

In this paper we are going to discuss how the adoption of a blockchain-based surveillance system can achieve all the desiderata listed above, while being a robust, transparent and cheap solution.

## Blockchain Architecture

Blockchain systems can vary in their specifications. But there are a few basic features which are key to the solution we propose here. Mainly they must be decentralized, which rules out “permissioned” blockchain systems, and support the implementation of smart-contracts, which consist autoenforced logic without centralized control. For this work we will assume Ethereum(Wood, 2014) as the underlying blockchain but we must keep in mind that this solution can easily be adapted to other platforms.

Let’s start with some definitions. Let us call a *user*, each reporter in the surveillance network, it can be a health professional, a clinic or a hospital. Users connect to the blockchain via client software which can emit proper transactions to be recorded onto the blockchain.

Each *user* will have an *address* in the Ethereum address space, which works as an unique identifier of the *user* and this address will be part of each and every transaction submitted by the *user*.

Let a *transaction* mean a case report broadcast to the Ethereum network via the client software.

A *private key* is associated to each address, and can be used to *sign* a *transaction*. Each transaction is signed and timestamped before being broadcast to the network.

Before a transaction can be recorded on the blockchain, it must satisfy the logic of the smart-contract controlling the surveillance system, but also have its signature validated, making it impossible to impersonate a user without access to its *private key*. Once transactions get recorded on the blockchain they are permanent making it impossible to erase data. Since the entire blockchain is also replicate on every node of the ethereum blockchain, the possibility of data loss is negligible.

The smart-contract governing the rules applicable to the surveillance workflow will not be detailed here since it will vary according to the application, but it suffices to say that its logic should focus sctrictly on the required validation of case reports, leaving higher order management decisions to humans.

### Case Reports Data Structure and Blockchain Publishing

Typically it is not feasible to store data within a blockchain as it would drive the cost of transactions to an ureasonably high value. However, there are alternatives for decentralized storage of data records of arbitrary size, such Ethereum’s Swarm and IPFS.

Therefore we propose that case reports generate two different records: The actual data from the case report forms wich will be stored on IPFS or Swarm, and its cryptographic hash will be stored in the blockchain as explained below.

We can obtain the cryptographic hash of a case report record by loadng it into a data structure know as a Merkle tree (figure 3). In a Merkle tree the data is a tree where each node is a hash of the sum of of their children node’s hashes. The root of the tree serves as an unique id for the entire record, since any other transaction differing on at least one node will yield a different root hash. Only the Merkle tree root hash will then be store into the blockchain to serve as a permanent record that that exact report was created at that timestamp by the reporter who signed the transaction(figure 2).

**Figure 1:**
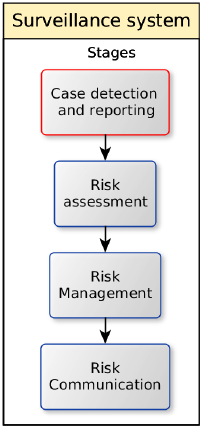
Structure of a surveillance system. We are concerned only with the red block.

**Figure 2:**
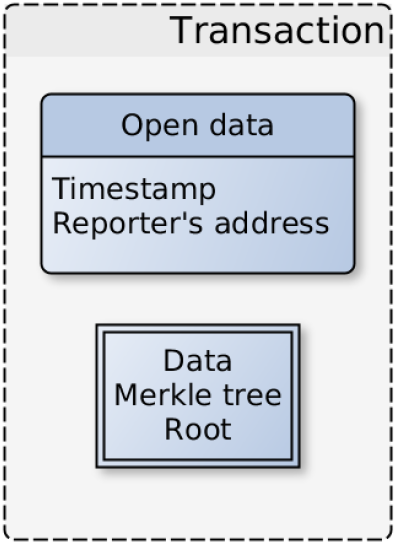
Contents of a transaction. Standard Ethereum transaction plus data tree root hash

**Figure 3:**
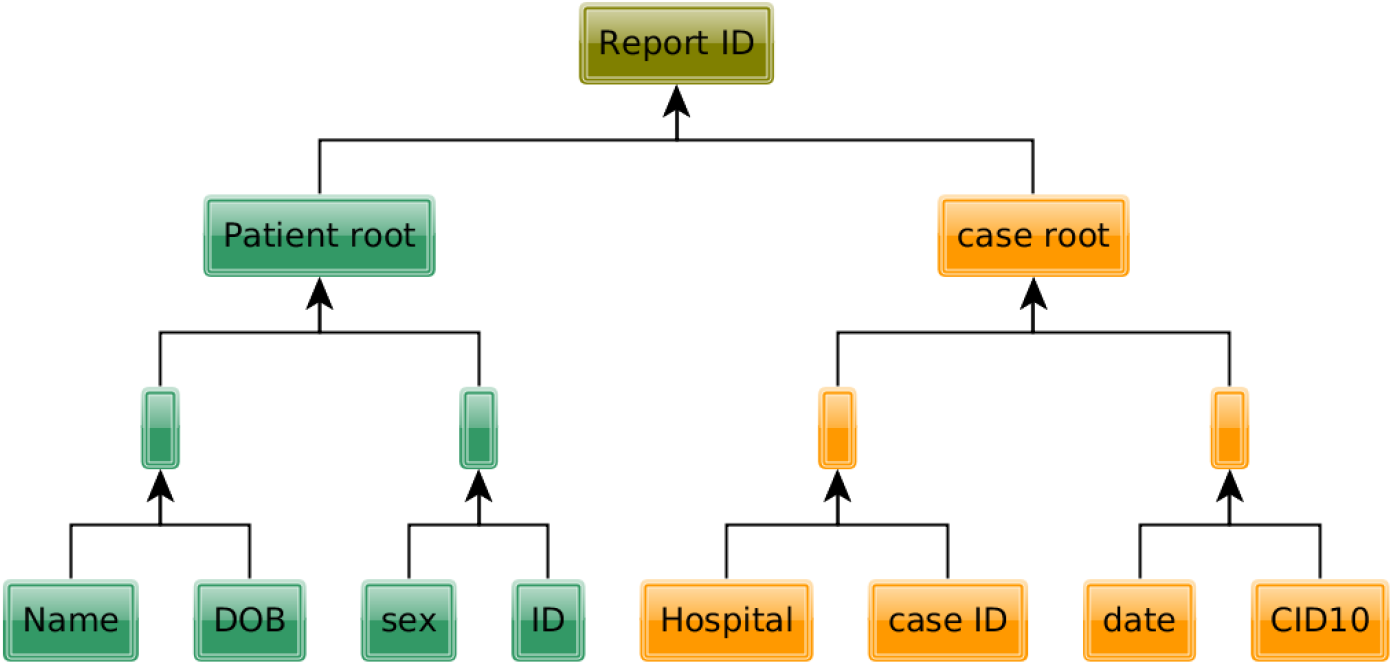
Merkle tree containing the case report data. This is a toy data model, real case reports may have as many fields as required.

Its interesting to notice that if we group the leaves (data fields) by category we can have branch roots which act as category tags or IDs. In the example of figure 3, we select all reports from the same patient by filtering on the patient root. Besides the patient and case roots shown in the example tree, we can have a location root, a clinical symptoms root etc. defined according to the reporting protocol. These roots can be used to filter or group cases based on any of the categories they represent.

Now we are going to show how the features of the blockchain, can help us achieve the desiderata elicited before.

### Provenance

As stated before, transactions originate from users, users are associated to blockchain addresses and each address to a private key. This private key is used to sign all transactions (case reports), making it impossible (short of stealing the private keys) for anyone to impersonate a user to create fake reports. The signature checks are a built-in feature of the blockchain and are performed by all blockchain nodes replicating the transactions. The records sotored in the IPFS or SWARM may also include the blockchain transaction ID making them permanently linked.

### Timing

Reports are timestamped. So the exact time of the report is registered automatically and cannot be manipulated by the node. Timestamps are used in the validation algorithm. Accurate timing does not imply synchronous operation, on the contrary since the timestamp is generated locally in the blockchain client, it allows for asynchronous publication without loss of the proper temporal ordering of events.

### Uniqueness

Reports are born unique due to the combination of the timestamp, user address and other data from the report. The Merkle tree root hash of these fields is inserted into the data field of a standard blockchain transaction which also has an unique transaction ID.

### Selective Privacy

Since we are proposing recording the data on a plublic decentralized data store, the case report data payload, contains all PII in encrypted form. Information such as timestamp, reporting establishment ID and other pieces of information deemed public in compliance with local legislation, can remain unencrypted and visible to anyone.

### Queryability

Due to the mixed open/encrypted composition of the data. The records allow for extensive SQL-like queries. Open fields can be fully queried, and encrypted fields although not adequate to filtering, can be used for groupings, for example one can find the number of cases per household without knowing where the households are located.

### Coverage

Traditional information systems rely of fixed computer terminals and local network connections. The blockchain client nodes can be packed in a simple mobile App, it can be deployed at next to zero cost. Since it does not require centralized supervision, it can scale without limits even when to places where connectivity is deficient, since constant connectivity is not required.

### Incentives

All collaborative systems require some form of incentive to work. In this case, a smart-contract associated with the DAG or Blockchain will emit tokens in exchange for the validation of reports(transactions). the basic validation algorithm for a report to be included in the blockchain is fixed. But other higher level validation tasks can be defined and rewarded via the smart-contract.

### Consensus

Reports of the same disease for the same patient at different places at roughly the same time can be detected and pruned according to some criterion. This type of validation can be done by epidemiologists rewarded by tokens.

### Versioning

Updates to reports which have already been reported can be registered into the blockchain as a new report with a reference to the unique ID of the original report thus the full history of the case gets preserved. When querying for cases, updated reports are clearly identifiable and can be used instead of the original reports.

### Durability

The records stored on the blockchain are permanent, and replicated on a large number of nodes making them robust to data corruption or loss.

## Deployment

The implementation and deployment of this distributed-ledger-based surveillance system can rely on different blockchain platforms. Even though we presented the solution assuming Ethereum blockchain, The only requirement is that the platform allows for a data field in the transaction data structure which can accomodate 32 bytes (256 bit hash) corresponding to the data tree root hash.

Eventhough it can be useful, smart-contract support is not required. A minimal version of the solution proposed here can work based on simple blockchain transactions and a standard database management system for the data storage. The alternative decentralized storage layer proposed, IPFS (Benet, 2014), can work independently of the blockchain layer chosen.

## Discussion and Conclusion

Adopting a distributed ledger to record disease case reports can bring several advantages over the current information systems backing disease surveillance. Among the main advantages is the immediate validation and availability of data, which can lead to faster responses of health monitoring systems during public health emergencies.

The distributed storage of the database offers greater transparency, through data locality, and also eliminates the problem of having central servers as a single point of failure. This means that reporting will never be delayed by system failures and is always accessible for reads.

The existence of a blockchain-based surveillance system such as the one described here is not incompatible with the maintenance of the old system, since we can have reports being sent simultaneously to the two databases. But we believe that gradually the distributed surveillance system will make the traditional information system obsolete.

The integration of the surveillance system with smart-contracts opens new possibilities in the form of a rich system of incentives for data validation and analysis.

We have only looked at the optimization of the first stage of a disease surveillance system through the adoption of distributed ledger technology. However once this stage is on a blockchain, other possibilities open-up, for example a marketplace for analytical models could be built on top of these open records. Any predictive model in this marketplace would be automatically comparable since they are using identical data. If the source code for the models is stored in public repositories they can be easily updated whenever the data changes. Moreover, Public Health agencies could fund the development of analytical models, directly through smart-contracts which would control the validation of the results releasing payments as the research project achieves pre-determined milestones, which can be validated automatically.

In summary, there is great benefit for a secure continuous release of data. The benefits of open public data are not a new idea, but we believe that distributed ledger technology is the key to open sensitive datasets in an effective and secure way.

## Footnotes

1 Personally identifiable information

